# Efficacy and immunogenicity of different BCG doses in BALB/c and CB6F1 mice when challenged with H37Rv or Beijing HN878

**DOI:** 10.1101/2020.10.21.349373

**Authors:** Bhagwati Khatri, James Keeble, Belinda Dagg, Daryan A. Kaveh, Philip J. Hogarth, Mei Mei Ho

## Abstract

In this study, 2 strains of mice (BALB/c and CB6F1) were vaccinated with a range of Bacille Calmette-Guérin (BCG) Danish doses from 3×10^5^ to 30 CFU/mouse, followed by either immunogenicity evaluation or aerosol infection with *Mycobacterium tuberculosis* (a laboratory strain H37Rv or West-Beijing HN878 strain). The results indicated that both strains of mice when infected with HN878 exhibited significant protection in their lungs with BCG doses at 3×10^5^ – 3000 CFU (BALB/c) and 3×10^5^-300 CFU (CB6F1). Whereas, both strains of mice when infected with H37Rv, significant protection was seen in BCG doses at 3×10^5^ - 300 CFU. Immunological evaluation revealed interesting results; i) both strains of mice demonstrated a significant increase in the frequencies of BCG-specific IFNγ^+^ IL2^+^ TNFα^+^ CD4 T cells in the BCG doses at 3×10^5^ – 3000 CFU (BALB/c) and 3×10^5^ - 300 CFU (CB6F1); ii) secretion of IL2 and IFNγ were correlated with the bacterial burden in the lungs of HN878 infected CB6F1 mice. The study demonstrated a BCG dose at 3000 CFU (an equivalent single human dose in the mice by body weight index) is protective in both strains of mice and the use of a virulent clinical isolate in testing new tuberculosis vaccine/advancing research is recommended.

## Introduction

Bacille Calmette-Guérin (BCG) is one of the most widely used injectable vaccines, administered predominantly in neonates, but confers incomplete and variable protection against pulmonary tuberculosis (TB) in humans ^1, 2, 3^. Despite its widespread use, an estimated 1.5 million people in 2018 died due to the TB disease ^4^. Studies in the field of infectious diseases, especially for intracellular pathogens, acknowledges the complex interplay of the magnitude, speed and qualitative nature of protective immune responses by chronic infections (such as TB, leprosy, leishmaniasis) which is not as straightforward as an effective antibody response against extracellular pathogens ^5^.

Although there are many animal models which are used in conducting TB research and testing vaccine candidates or drugs against TB, the mouse model is the most preferred choice mainly due to the small animal size, vast array of commercially available reagents and low cost of running an experiment. Having recognised the limitations of animal models in terms of predicting outcomes in humans, moving forward, it is vital to improve animal models wherever possible ^6, 7^. From the decades of published animal TB challenge studies ^7, 8, 9, 10^, it is evident to state that protective responses induced by BCG vaccine against TB in mice varies when compounded by the choice of mouse strain ^11^, strains of *Mycobacteria tuberculosis (Mtb)* for infection, route of immunisation and the interval between BCG vaccination and *Mtb* challenge. Therefore, our aim was to optimise some of these parameters by analysing the efficacy of different doses of BCG against two strains of *Mtb* namely, laboratory TB strain H37Rv and a virulent clinical strain, W-Beijing HN878 (HN878) and using each strain of *Mtb* in the head-to-head comparison using two types of mouse strains, BALB/c and CB6F1 (F1).

The study was divided into two parts; 1. H37Rv or HN878 challenge - different doses of BCG were evaluated by measuring *Mtb* burden in the lungs and spleen of mice infected with either *Mtb* strain and; 2. Immunogenicity evaluation - cellular immune responses were analysed using flow cytometry, IFN-γ ELISPOT and multiplex cytokine/chemokine assays in equivalent BCG vaccinated and control mice.

The overall results demonstrated differences in the protection and cellular immune responses in both strains of mice. Our findings will benefit refinement of the TB challenge mouse model for the pre-clinical testing and selection of new TB vaccines.

## Results

In the initial experiment where BALB/c and F1 mice were vaccinated with a full range of BCG doses (3×10^5^, 3×10^4^, 3000, 300 or 30 CFU/mouse) followed by H37Rv challenge, a comparable protection was seen in the lungs of mice vaccinated with BCG dose at 3×10^4^ and 3000 CFU (Fig. 1A). Considering the principle of replacement, reduction and refinement (3Rs) in experimental design using animals, BCG dose at 3×10^4^ CFU was omitted from HN878 challenge experiments and the immunogenicity evaluation study. Concentrations of all serially diluted BCG doses (CFU/ml) were plated on to 7H11 agar plates and confirmed to be within the accepted range (Supplementary Table 1). An infection inoculum for H37Rv or HN878 were also confirmed using 7H11 agar plates and shown to be within the expected range of 3.6-5 x 10^6^ CFU/ml or 4-7 x 10^6^ CFU/ml, respectively to achieve low dose aerosol infection of approximately 100 CFU per mouse. The result for low dose infection in the lungs of BALB/c mice were, H37Rv - average 71 ± 63 (Std. Dev) CFU and HN878 – average 144 ± 64 (Std. Dev) CFU per mouse (Supplementary Table 2).

**Figure 1:**
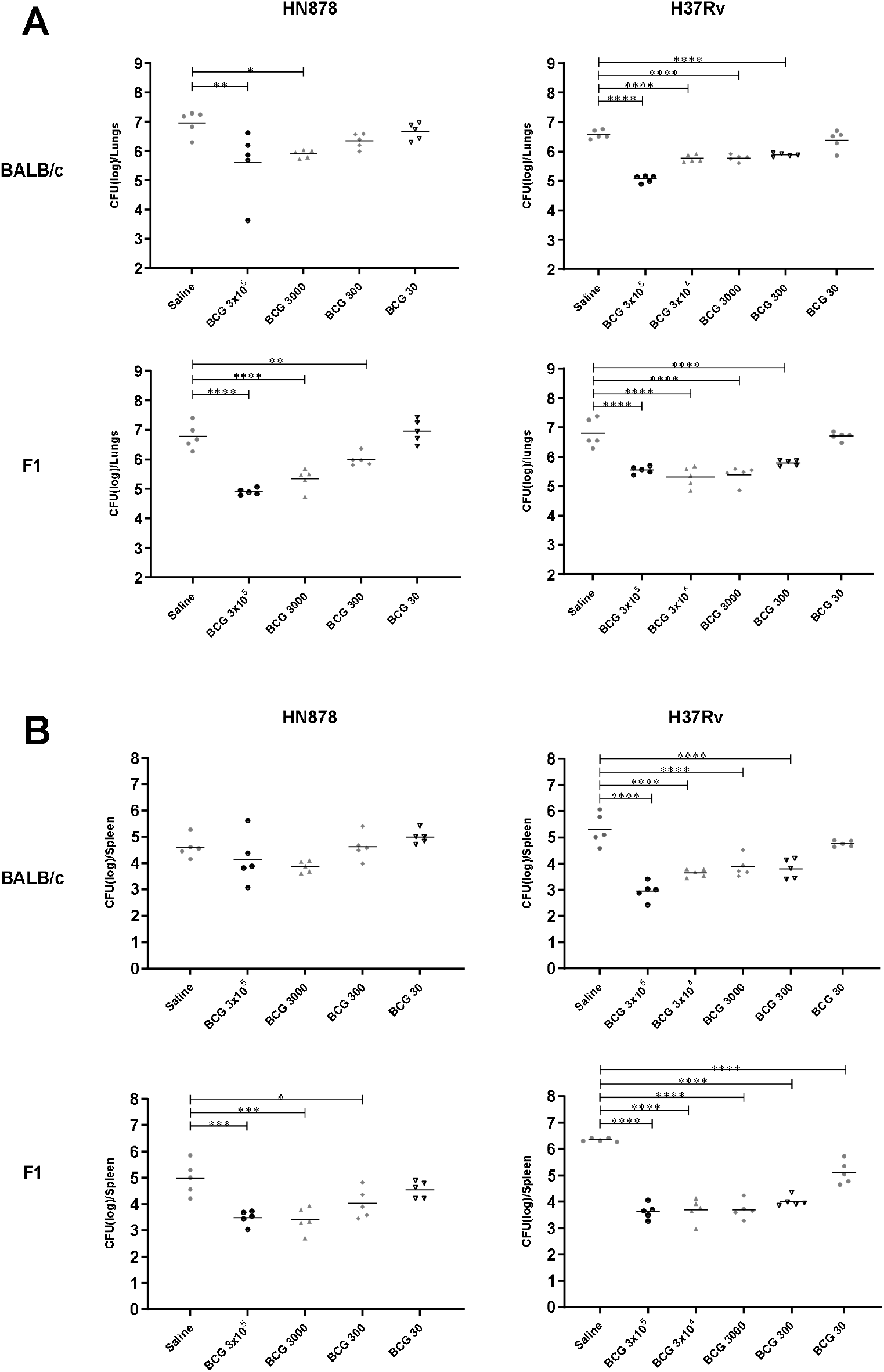
The CFU data for BALB/c and F1 mice vaccinated with various BCG doses and challenged with HN878 and H37Rv. Four weeks post infection, lungs and spleen were harvested, and bacterial loads were determined. A, Lungs CFU data as Log_10_ protection for the vaccinated BALB/c and F1 mice. B, Spleen CFU data as Log_10_ protection for the vaccinated BALB/c and F1 mice. One-way ANOVA, Tukey’s multiple comparison test was performed for BALB/c and F1 mice infected with H37Rv and HN878 strains. *p<0.05, **p<0.005, ***p<0.0005, ****p<0.0001.

### BCG protection in BALB/c and F1 mice varies when challenged with H37Rv or HN878

Four weeks after BCG vaccination, BALB/c and F1 mice were infected with either H37Rv or HN878. Animals were sacrificed at 4 weeks post infection and the lungs and spleen were harvested to determine *Mtb* burden. Fig. 1A shows the graphical representation of the protection data expressed as Log_10_ CFU in the lungs and spleen of BALB/c and F1 mice infected with either H37Rv or HN878. Tables 1A and 1B represent the differences in the mean of Log_10_ CFU/organ between all the BCG vaccinated and the control groups, including within BCG vaccinated groups in the lungs and spleen of both strains of mice when infected with H37Rv or HN878. For the lowest BCG dose at 30 CFU, *Mtb* burden (Log_10_ CFU/organ) in the lungs of both mouse strains infected with either H37Rv or HN878 were equivalent to the control group (Fig. 1A; Table 1A,1B).

**Table 1:**
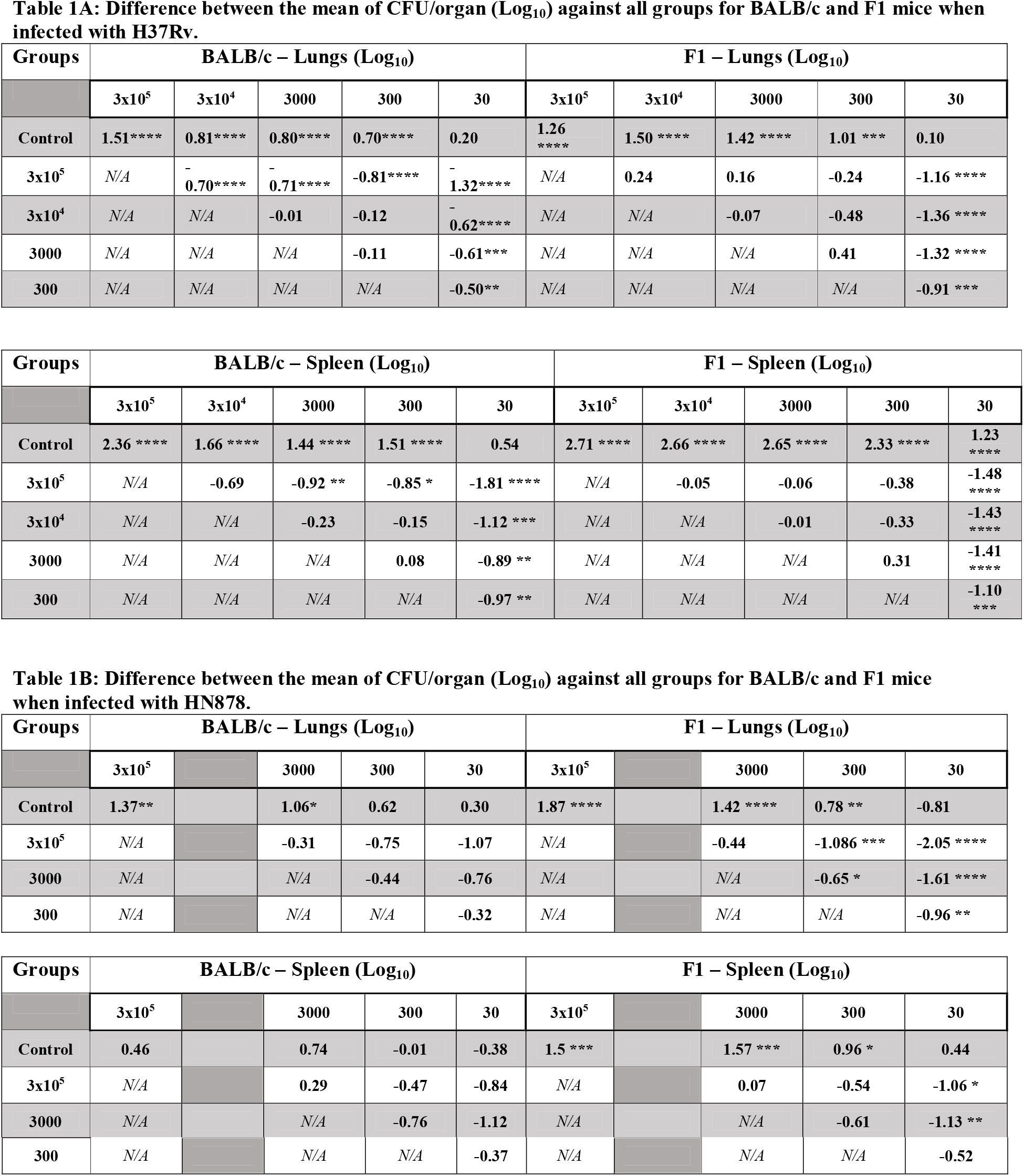
Lung and spleen protection data for BALB/c and F1 mice when infected with H37Rv or HN878 strains. The values in the Log_10_ protection data is the difference in the means of control group when compared to the BCG dose groups and the difference within BCG groups. Data is the representation of two independent experiments for H37Rv and one experiment for HN878. Values representing difference for the mean Log_10_ of the BCG and control groups (left column) – mean Log_10_ of the BCG groups (Top Row). A, Difference between the means of the lung and spleen of Log_10_ values for BALB/c and F1 mice infected with H37Rv. B, Difference between the means of the lung and spleen of Log_10_ values for BALB/c and F1 mice infected with HN878. One-way ANOVA, Tukey’s multiple comparison test was performed for BALB/c and F1 mice infected with H37Rv or HN878. *p<0.05, **p<0.005, ***p<0.0005, ****p<0.0001.

H37Rv infected BALB/c and F1 mice vaccinated with BCG at 3×10^5^ - 300 CFU, displayed significant protection (Fig. 1A and Table 1) in the lungs when compared to the control group. The protection within the BCG groups for BALB/c and F1 mice differed. BALB/c mice infected with H37Rv showed significantly more protection in the lungs of the highest BCG dose 3×10^5^ CFU (Table 1A) when compared to all the lower doses of BCG: 3×10^4^ CFU (−0.7 Log_10_ CFU/lungs), 3000 CFU (−0.71 Log_10_ CFU/lungs), 300 CFU (−0.81 Log_10_ CFU/lungs) and 30 CFU (−1.32 Log_10_ CFU/lungs). In comparison to BALB/c, H37Rv infected F1 mice displayed comparable protection in the lungs with the highest dose of BCG 3×10^5^ CFU when compared to all the lower doses of BCG 3×10^4^ CFU (0.24 Log10 CFU/lungs), 3000 CFU (0.16 Log10 CFU/lungs), 300 CFU (−0.24 Log10 CFU/lungs), except 30 CFU (−1.16 Log_10_ CFU/lungs) (Table 1A). In the spleen (Fig. 1B), both strains of mice showed significantly more protection in the groups vaccinated with BCG at 3×10^5^ - 300 CFU when compared to the control group. In addition, spleen of F1, but not BALB/c mice, BCG dose at 30 CFU showed significant protection when compared to the control group, Fig. 1B and Table 1A (1.23 Log_10_ CFU/spleen).

When infected with HN878, BALB/c mice only demonstrated significant protection in the lungs with BCG doses at 3×10^5^ and 3000 CFU while, F1 mice showed significant protection in the lungs with a BCG dose as low as 300 CFU when compared to the control group (Fig. 1A and Table 1B). In addition, F1 mice displayed a dose response pattern in the protection (1.87, 1.42 and 0.78 Log_10_ CFU/lungs, respectively) when compared to the control group (Fig. 1A, Table 1B). In the spleen (Fig. 1B and Table 1B), F1 mice showed significant protection with BCG at 3×10^5^ - 300 CFU whereas, BALB/c mice failed to exhibit protection for all the BCG vaccinated groups when compared to the control group.

### IFNγ secreting cells by ELISPOT assay

Given the established critical role of IFNγ in controlling TB infection ^12, 13^, the frequencies of antigen-specific IFNγ secreting cells in ex vivo isolated cells from the spleen and the lungs of all mice were evaluated by ELISPOT. All splenocytes and lung cells were stimulated with a defined M7 protein cocktail. In the spleen and the lungs of BALB/c mice (Fig. 2A), a significantly high frequency of BCG-specific IFNγ secreting cells was evident in the mice vaccinated with BCG at 3×10^5^ and 3000 CFU whereas, they were equivalent to the control group in the BCG at 300 and 30 CFU groups. In the spleen of F1 mice, albeit the frequencies of IFNγ secreting cells in all the BCG vaccinated groups were not significantly different, BCG at 3×10^5^ – 30 CFU displayed higher frequencies of these cells when compared to the control group. However, in the lungs of F1 mice, only BCG at 3×10^5^ CFU found to have significantly higher frequency of IFNγ secreting cells whereas, BCG at 3000 and 300 CFU displayed higher responses compared to the control group.

**Figure 2:**
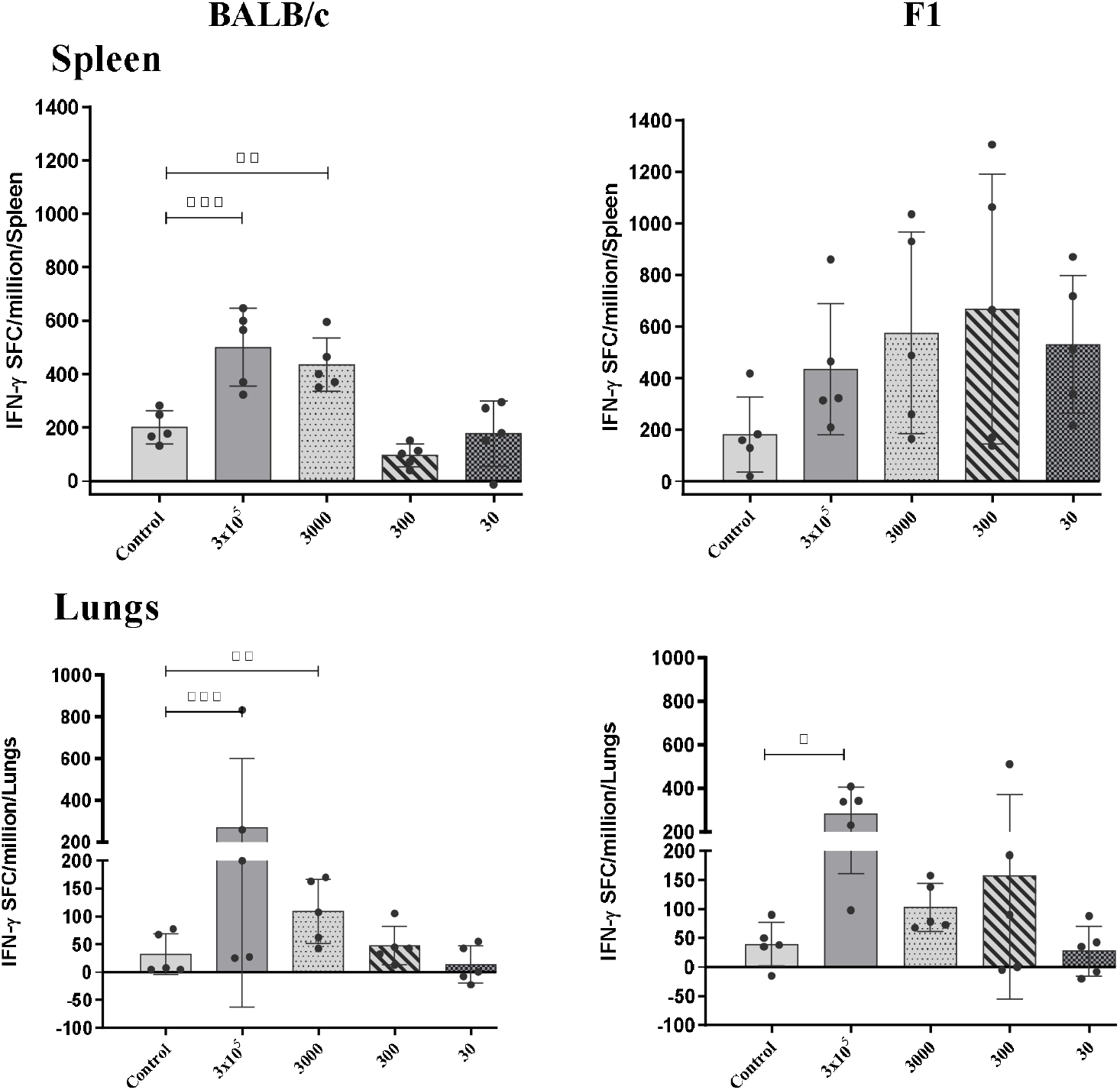
IFN-γ secreting cells in spleen and lungs of BALB/c and F1 mice. Splenocytes and lung cells isolated from six weeks post BCG immunized BALB/c and F1 mice and the frequency of IFN-γ secreting cells evaluated by *ex-vivo* ELISPOT. Approximately 2×10^5^ cells were cultured with M7 protein cocktail and developed by ELISPOT assay, bars representing mean (±S.E.) Spot Forming Cells (SFC) per million. Statistical analysis: One-way ANOVA, Dunnett’s test was performed. *p<0.05, **p<0.005, ***p<0.0005.

Together, these results highlight that in the spleen and the lungs of BALB/c and F1 mice, significant frequencies of BCG-specific IFNγ secreting cells were only detected in the groups that received either BCG at 3×10^5^ or 3000 (BALB/c mice only) and not the lower doses.

### Multifunctional CD4 T cells producing IFNγ, IL2 and TNFα

In order to better resolve the functionality of the responding T cells induced by BCG vaccination, M7 stimulated splenocytes from all groups of BALB/c and F1 mice were subsequently interrogated by intracellular staining (ICS) using 11 colour flow cytometric analysis. The frequency of CD4 T cells producing any combination of IFNγ, IL2, TNFα and IL17a production were assessed. The production of IL17a was not observed hence, omitted from the results for clarity. As shown in Fig. 3A, BALB/c and F1 mice induced significantly more multifunctional IFNγ^+^ IL2^+^TNFα^+^ CD4 T cells with BCG at 3×10^5^-3000 CFU and 3×10^5^-300 CFU, respectively when compared to the control group. Within the BCG vaccinated groups, 3×10^5^ CFU induced a significantly higher frequency of multifunctional IFNγ^+^IL2^+^TNFα^+^ CD4 T cells in BALB/c when compared to 3000 CFU and in F1 mice when compared to 3000 and 300 CFU.

**Figure 3:**
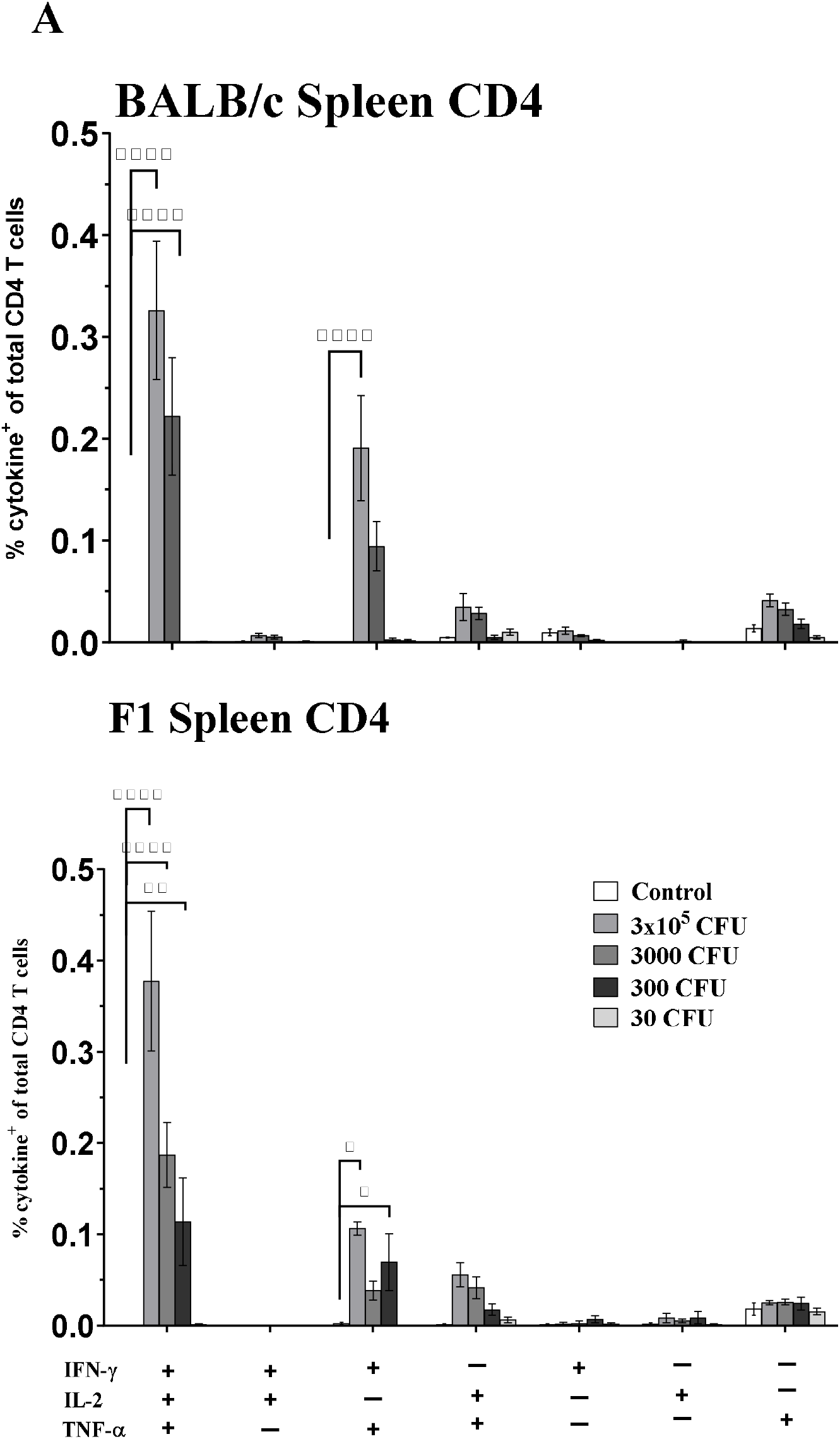
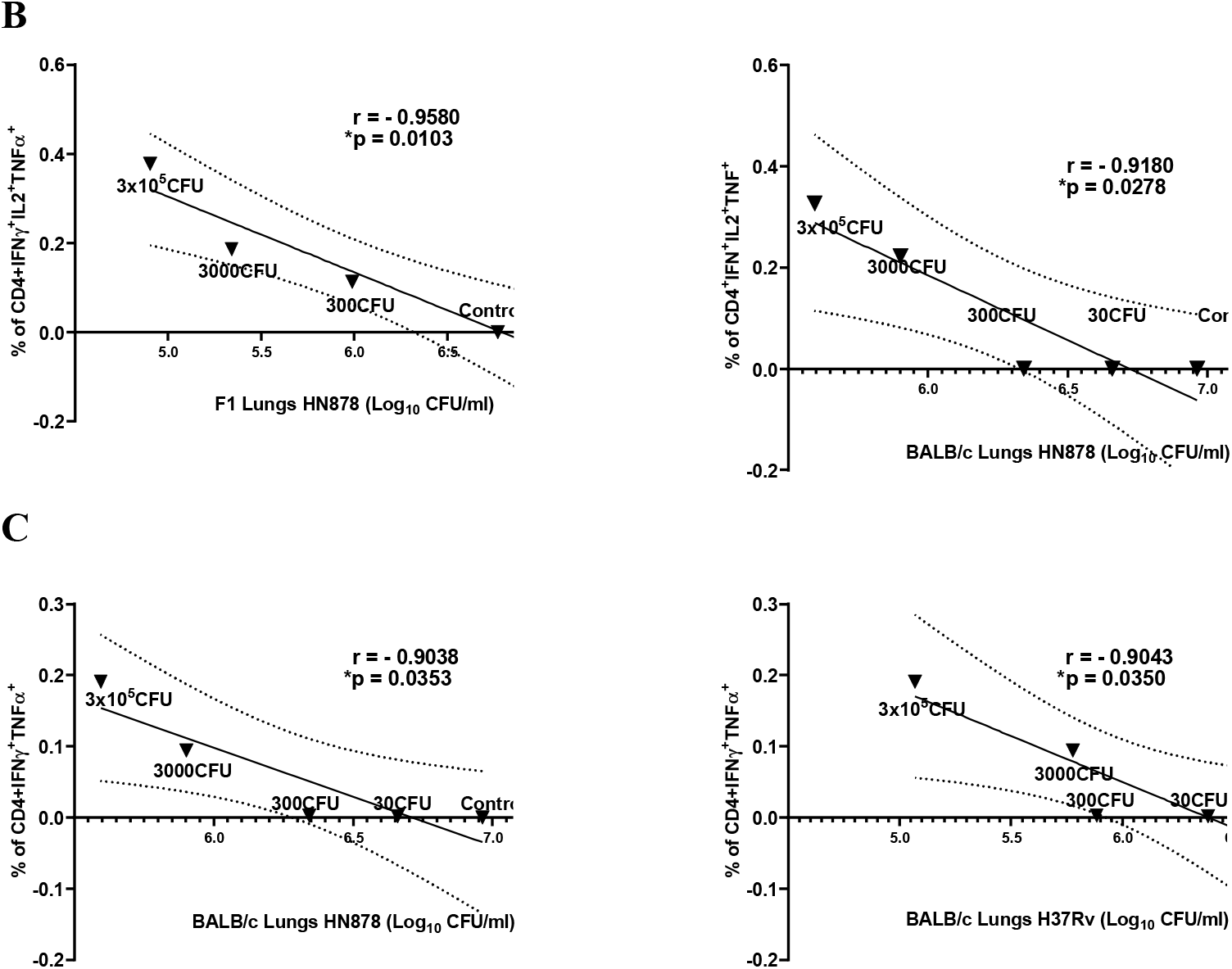
Vaccination induced resident multifunctional CD4^+^ cells in the spleen of BALB/c and F1 mice. The frequency of multifunctional CD4^+^CD44^hi^CD62L^lo^ cells at six weeks following BCG immunization. A, splenocytes from BALB/c and F1 mice for control, BCG groups - 3×10^5^, 3000, 300 and 30 CFU were isolated, stimulated with M7 proteins cocktail and stained by ICS. Bars represent mean (± SE) % frequency of cells of indicated T cell phenotype as a % of total CD4+ cells. Two-way ANOVA, Tukey’s multiple comparison was performed, *p<0.05, **p<0.005, ***p<0.0005, ****p<0.0001. B and C, correlation between the means of percentages of CD4^+^IFNγ^+^IL2^+^TNFa^+^ (B) and CD4^+^IFNγ^+^TNFα^+^ (C) for BCG groups (3×10^5^ – 30 CFU annotated onto the graphs) and control vs TB burden in the lungs of BALB/c and F1 mice expressed as Log_10_ CFU/lungs (as shown in Fig. 1A). Correlation coefficient was assessed using Pearson’s two-tailed correlation test with 95% confidence interval shown as dotted lines on B and C graphs

For both strains of mice, cytokine producing splenocytes IFNγ^+^ TNFα^+^ CD4 T cells were significantly induced in BCG at 3×10^5^-3000 CFU when compared to the control group (Fig. 3A). In addition, F1 mice induced IFNγ^+^ TNFα^+^ CD4 T cells in 300 CFU compared to the control group (Fig. 3A). None of the mouse strain exhibited multifunctional IFNγ^+^IL2^+^TNFα^+^ or IFNγ^+^ TNFα^+^ CD4 T cells in the BCG at 30 CFU group. The cytokine producing CD4 T cells detected here, displayed a CD44^hi^CD62L^lo^ phenotype, indicative of CD4 T Effector Memory (TEM), data not shown.

To understand the significance of correlation between the means of BCG-specific multifunctional CD4 TEM cells in the spleen and *Mtb* burden in the lungs of H37Rv or HN878 infected BALB/c and F1 mice in the equivalent vaccine groups, we performed correlation coefficient analysis. As shown in Fig. 3B and 3C, the correlations were analysed between the mean of IFNγ^+^IL2^+^TNFα^+^ or IFNγ^+^ TNFα^+^ CD4 T cells and the mean of *Mtb* burden as expressed in Log_10_ CFU/lungs. Multifunctional IFNγ^+^IL2^+^TNFα^+^ CD4 T cells were significantly inversely correlated with HN878 infected BALB/c (r = −0.918, *p* = 0.0278) and F1 mice (r = −0.958, *p* = 0.0103). However, correlation was not seen with H37Rv infected BALB/c (r = −0.8325, *p* = 0.0802) and F1 mice (r = −0.7844, *p* = 0.1162), data not shown. For IFNγ^+^ TNFα^+^ CD4 T cells, a significantly inverse correlation was exhibited in BALB/c mice infected with either H37Rv (r = −0.9038, *p* = 0.0353) or HN878 (r = −0.9043, *p* = 0.035). Again, for IFNγ^+^ TNFα^+^ CD4 T cells, correlation was not seen in F1 mice infected with H37Rv (r = −0.7844, *p* = 0.1162) or HN878 (r = −0.8641, *p* = 0.0589), data not shown.

These results highlight the induction of multifunctional IFNγ^+^IL2^+^TNFα^+^ or IFNγ^+^ TNFα^+^ CD4 T cells in both strains of mice exhibiting dose response to the vaccine. The highest BCG dose group, 3×10^5^ CFU induced significantly higher frequencies of these CD4 TEM cells when compared to the BCG at 3000 – 30 CFU groups. In addition, the IFNγ^+^IL2^+^TNFα^+^ CD4 T cells inversely correlated with the HN878 infected BALB/c and F1 mice which demonstrates, the higher the proportion of these cells, the lower is *Mtb* burden in the lungs of subsequently infected mice. Interestingly, IFNγ^+^ TNFα^+^ CD4 T cells inversely correlated with the *Mtb* burden in the BALB/c mice with both *Mtb* challenge strains, but not with F1 mice.

### Differential expression of cytokines/chemokines in BALB/c and F1 mice

Splenocytes and lung cells isolated from the BCG vaccinated and control groups were stimulated with M7 protein cocktail for 3 days and the supernatants were collected. Analyses of cytokines and chemokines, as listed in the Table 2, were performed in the supernatants of cultured cells (stimulated and unstimulated) using Bio-Plex Pro Mouse Chemokine Panel. For the data analysis, an estimated concentration (pg/ml) of cytokine/chemokine were interpolated from the linear part of the 5-parameter standard curve. Furthermore, the interpolated concentrations from the supernatants of stimulated cells were subtracted with those of unstimulated cells supernatant and these concentrations were used for the data analysis. Fig. 4A are the representative graphs of cytokines/chemokines obtained from the spleen or the lungs of BALB/c or F1 mice and, the concentrations for all cytokines/chemokines from different samples are presented as the radar plots with the numerical data tabulated in the supplementary Table 3. Groups vaccinated with BCG at 3×10^5^ CFU exhibited greater variability in the lungs of BALB/c mice compared to the other BCG vaccinated groups. For the ease of result presentation, significance in all the BCG vaccinated groups have been compared with the control group, unless depicted otherwise.

**Figure 4:**
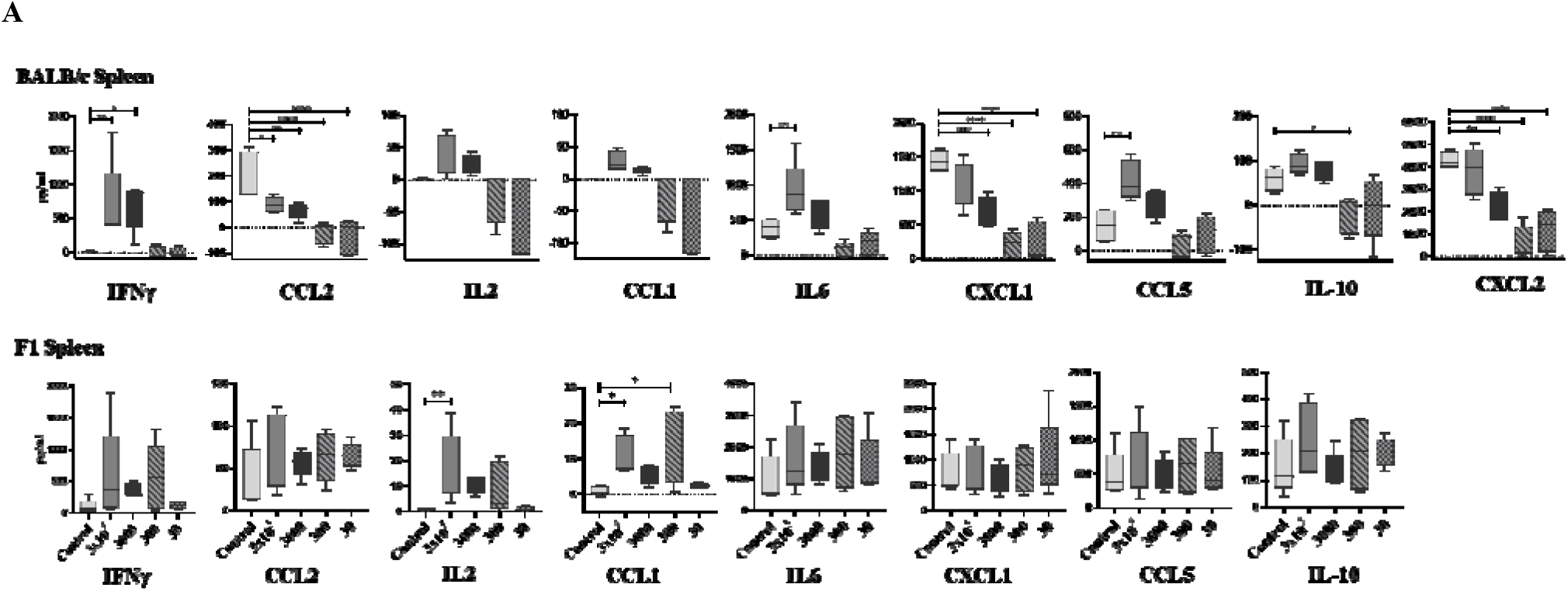
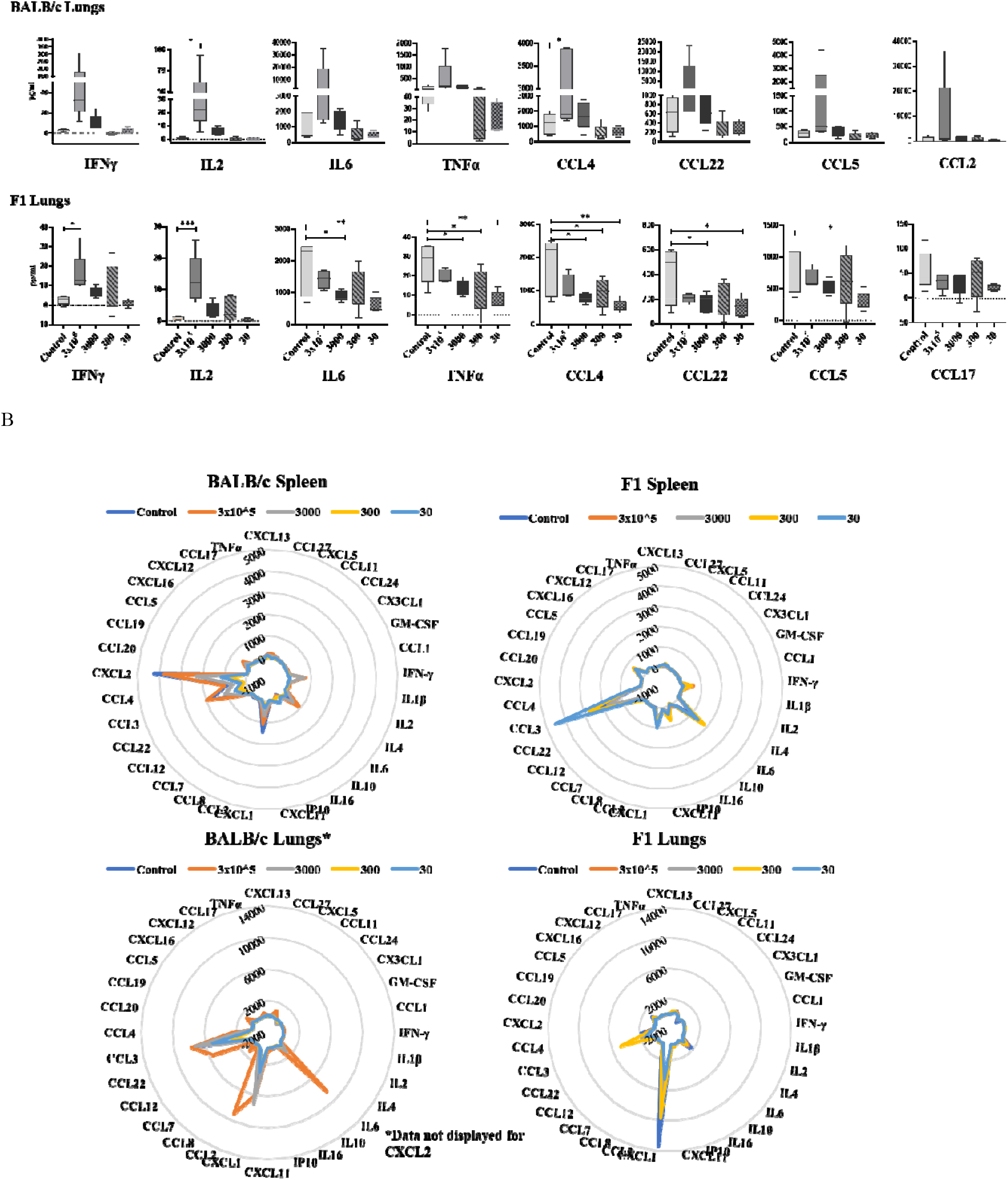
Cytokines/ chemokines expression in the M7 stimulated splenocytes and lung cells from BALB/c and F1 mice vaccinated with various BCG doses. A; representative graphs of cytokines/ chemokines expressed in pg/ml after 3 days stimulated splenocytes (top graphs) and lung cells (bottom graphs) of BALB/c and F1 mice for all groups. B; Radar or spider plot with cytokine/chemokine at the corners of the plot. The results of the mean concentration (n=5) for all cytokines/chemokines measured in the stimulated splenocytes and lung cells of BALB/c (top and bottom left graphs) and F1 (Top and bottom right graphs). One-way ANOVA, Dunnett’s multiple comparison test, *p<0.05, **p<0.005, ***p<0.0005 was performed for A and B.

**Table 2:** Classification of examined cytokines and chemokines in the M7 stimulated supernatants of splenocytes and lung cells from BCG vaccinated BALB/c and F1 mice.

|  |  |  |
| --- | --- | --- |
| **Chemokine** | **CCL** | CTACK/CCL27, Eotaxin/CCL11, Eotaxin-2/CCL24, I-309/CCL1, MCP-1/CCL2, MCP-2/CCL8*, MCP-3/CCL7, MCP-5/CCL12, MDC/CCL22, MIP-1α/CCL3, MIP-1β/CCL4, MIP-3α/CCL20, MIP-3β/CCL19, RANTES/CCL5, MCP-3/CCL7 and TECK/CCL25* |
|  | **CXCL** | BCA-1/CXCL13, Fractalkine/CX3CL1, ENA-78/CXCL5, I-TAC/CXCL11, KC/CXCL1, MIP-2/CXCL2*, SCYB16/CXCL16, SDF-1α/CXCL12, IP10/CXCL10 |
| **TNF** |  | TNFα |
| **Interleukins Type I** | Th1 cytokines growth factors and Interferons | IFNγ, IL2, IL16, GM-CSF, IL1β |
| **Interleukins Type II** | Interleukins | IL4, IL10, IL6 |

\*Not analysed for F1 mice (Splenocytes and lung cells) as absent in the Bio-Plex Pro Mouse Chemokine Panel.

As expected, secreted IFNγ concentration in the splenocytes of BALB/c mice (Fig. 4A) were detectable at significantly higher concentration in the BCG at 3×10^5^ and 3000 CFU groups. However, although IFNγ responses were not significantly different, the splenocytes of F1 mice displayed an elevated response in the BCG at 3×10^5^ – 300 CFU groups (Fig. 4A). Interestingly, a significantly higher IL2 concentration (Fig. 4A&B) in the BCG at 3×10^5^ CFU group for the F1 (spleen and lungs) and BALB/c (lungs) mice were observed. The elevated responses of IL2 were also detected in the splenocytes and the lung cells of F1 (BCG at 3000 - 300 CFU groups) and BALB/c (BCG at 3000 CFU group) mice.

Surprisingly, the concentrations of secreted chemokines, CCL2, CXCL1 and CXCL2 were significantly lower from the splenocytes of all the BCG vaccinated compared to the control group for BALB/c, but not F1 mice. For F1 mice, secreted IL6, TNFα, CCL4 and CCL22 concentrations from the lung cells were lower in all the BCG vaccinated groups and displaying significant differences in the BCG at 3000 – 30 CFU groups when compared to the control group. In contrast to the F1 mice, the lung cells of BALB/c mice secreted significantly higher concentrations of IL6 and CCL4 for 3×10^5^ CFU BCG group when compared to the control group. The rest of the cytokines/chemokines shown in Fig. 4A secreted elevated concentrations in the 3×10^5^ CFU BCG group when compared to the control and 3000 – 30 CFU BCG groups.

An overview of the graphical visualisation, all concentrations of cytokines/chemokines in the lungs and spleen of BALB/c and F1 mice have been presented in the radar plots (Fig. 4B) except, for CXCL2 from the supernatants of the lung cells of BALB/c mice which were substantially higher in concentration. Therefore, for the visual representation in the radar plot for BALB/c (lungs), CXCL2 concentration have been removed.

### Correlation of cytokine/chemokine with TB burden in the lungs of BALB/c and F1 mice

IFNγ is the only soluble immunological marker currently used to identify TB infection in the clinical settings by using an IFNγ release assay. Therefore, we sought to investigate the correlation of vaccine induced IFNγ and other cytokines/chemokines with the *Mtb* burden in the lungs of BALB/c and F1 mice subsequently infected with either H37Rv or HN878.

Firstly, correlation between the IFNγ concentration and other cytokine/chemokine concentrations from the stimulated cell supernatants were investigated (Supplementary Table 4). For the correlation analysis, the mean of IFNγ concentration was correlated with the mean concentration of other cytokines/chemokines from the supernatants of splenocytes/lung cells of BALB/c and F1 mice. For example, the mean of IFNγ concentration from BALB/c splenocytes was correlated with the mean concentrations of other cytokines/chemokines within the same supernatant sample. As shown in the Supplementary Table 4, IFNγ responses in the splenocytes of BALB/c mice significantly correlated with IL6 (p = 0.0406) and CCL5 (p = 0.0229). In the splenocytes of F1 mice, IL2 (p = 0.037), CCL1 (p = 0.0062) and CCL5 (p = 0.0441) were significantly correlated with the IFNγ responses. In addition, IL2 responses were strongly correlated with the IFNγ responses in the lung cells of BALB/c (p = 0.0002) and F1 (p = 0.0088) mice.

To gain further insight for whether the cytokines/chemokines which correlated with the IFNγ responses in the BCG vaccinated groups (Supplementary Table 4) also correlated with the *Mtb* burden in the lungs of BALB/c or F1 mice, therefore, we investigated using H37Rv or HN878 infected BALB/c and F1 mice (Fig. 5 and Table 3). The analysis produced an intriguing observation for IL2 correlation. Spleen IL2 (Fig. 5B, Table 3) responses were significantly inversely correlated with the lungs Log_10_ CFU of F1 mice infected with both strains of *Mtb* - H37Rv (p = 0.0310) or HN878 (p = 0.0032) whereas, BALB/c mice did not show any statistically significant correlation with IL2 responses. As expected, IFNγ responses in the lungs (Fig. 5A, Table 3), but not in the spleen (Fig. 5A) of F1 mice significantly correlated with the *Mtb* burden in the lungs of HN878 infected F1 mice. In contrast, only HN878 infected BALB/c mice showed significant correlation of *Mtb* burden in the lungs with the spleen IFNγ (Table 3).

**Figure 5:**
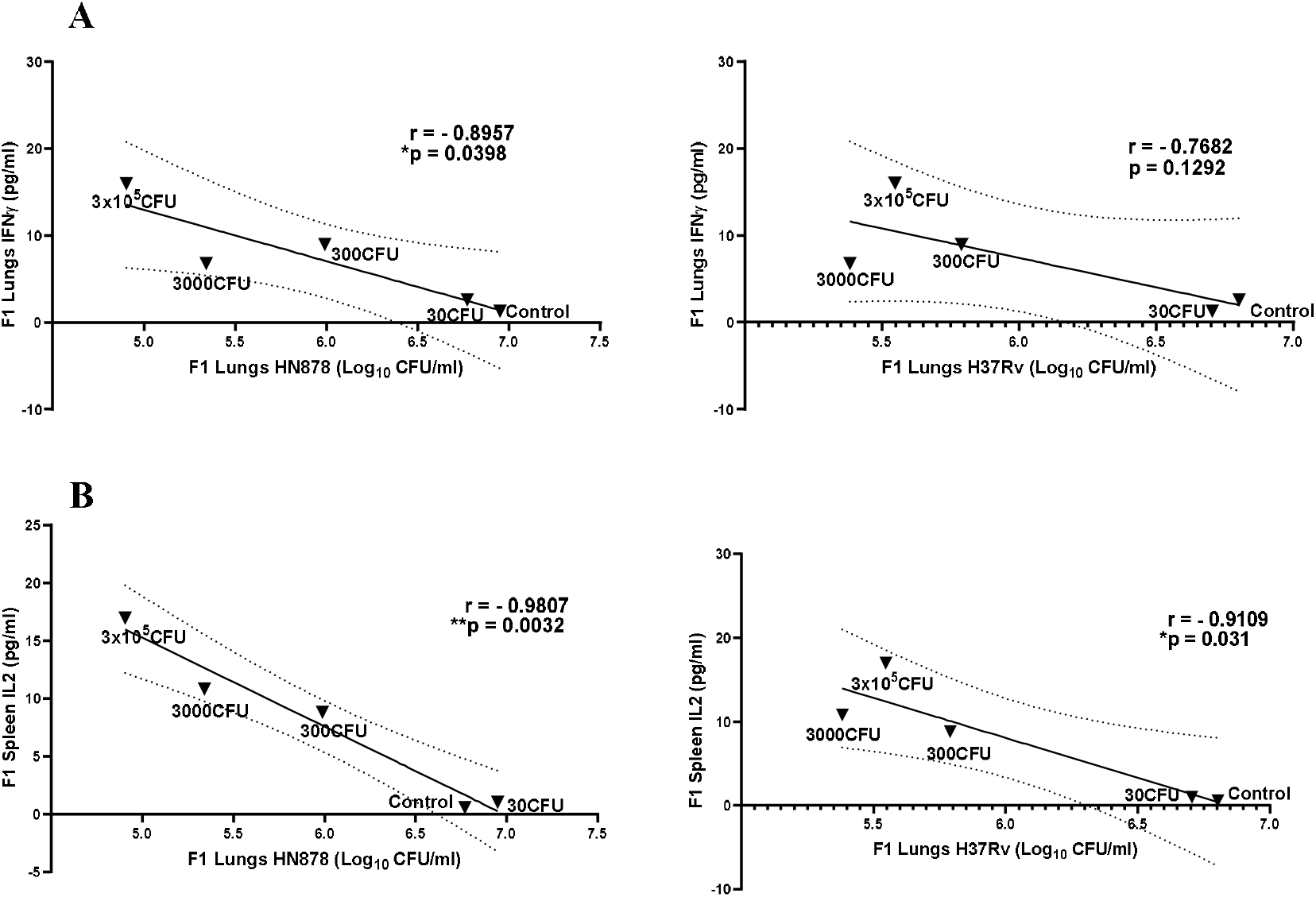
Correlation of cytokines from supernatants of M7 stimulated splenocytes and lung cells with TB burden in lungs of H37Rv or HN878 infected F1 mice. Representative graphs for correlation analysis between the means of cytokines from the supernatants of stimulated splenocytes/lung cells and TB burden (Log_10_ CFU/ml) in the lungs of H37Rv or HN878 infected F1 mice. Various doses of BCG vaccines have been annotated onto the graphs. A; The left and right graphs represent correlation between TB burden in the lungs of infected F1 mice (HN878 or H37Rv respectively) and IFNγ response in the supernatants of stimulated lung cells. B; The left and right graphs represent the correlation between the TB burden in the lungs of infected F1 mice (HN878 or H37Rv respectively) and IL2 responses in the supernatants of stimulated splenocytes. Correlation coefficient was assessed using Pearson’s two-tailed correlation test with 95% confidence interval shown as dotted lines in A and B

**Table 3:** Correlation between the cytokines from supernatants of M7 stimulated splenocytes or lung cells and *Mtb* burden in the lungs (Log_10_ CFU/lung) of H37Rv or HN878 infected BALB/c and F1 mice.

|  | **BALB/c H37Rv Lungs (Log<sub>10</sub> CFU/lungs)** | **BALB/c HN878 Lungs (Log<sub>10</sub> CFU/lungs)** | **F1 H37Rv Lungs (Log<sub>10</sub> CFU/lungs)** | **F1 HN878 Lungs (Log<sub>10</sub> CFU/lungs)** |  |
| --- | --- | --- | --- | --- | --- |
|  | - | **Spleen IFNγ** | **Spleen IL2** | **Lung IFNγ** | **Spleen IL2** |
| **Pearson r** | - | - 0.9033 | -0.9109 | - 0.8957 | -0.9807 |
| **P value (2-tailed)** | - | 0.0356 | 0.0310 | 0.0398 | 0.0032 |
| **Sig** | - | * | * | * | ** |
| **95% confidence intervals (Slope)** | - | -1176 to -80.07 | -17.46 to -1.595 | -11.32 to -0.5208 | -10.51 to -4.876 |

## Discussion

NIBSC have been supporting the international efforts to develop new TB vaccines using a specialised murine aerosol TB challenge model for testing new vaccine candidates. As a recognised WHO reference laboratory, NIBSC established the WHO reference reagents for BCG vaccine of various substrains ^14, 15^. Since then, our laboratory has been involved in investigating the mechanism of protective immune responses elicited by the BCG and a continuous effort in refining TB mouse models as the first line of screening tool for the selection of new TB vaccine candidates.

This study was designed to accommodate and address suppositions or observations as 1. To evaluate protection of various doses of BCG in BALB/c and F1 mice when infected with H37Rv or HN878 and to understand the protective cellular immunity induced by various BCG doses; 2. For the refinement of TB mouse model, we were interested to evaluate protective responses of BCG dose at 3000 CFU. BCG dosage at 1-4 x 10^5^ CFU is given to infants, preferably at the time of birth. The commonly used BCG dose in mice is also at 1-4 x 10^5^ CFU/mouse. If the body weight of an infant and an adult mouse is considered, the BCG dose at 3x 10^5^ CFU given to a mouse would be around 150 times in excess compared to the BCG dose in an infant. Therefore, most of the research conducted in mice using BCG dose at 1-4 x 10^5^ CFU would be around 2 Log higher compared to human and the immune parameters measured would fall under the high dose BCG responses. According to the study by Bretscher ^16, 17^ high dose of BCG would tend to skew the immune response more towards the mixture of Th1 and Th2. Therefore, it was pertinent to evaluate the protection and immune responses of 3000 CFU in mice, an equivalent dose of BCG 3 x 10^5^ CFU in infants; 3. For the BCG dose sparing in preclinical studies, concerns were raised in 2016 ^18^ for the shortage of global BCG vaccine and the published study in 2019 ^19^ highlighted the effect of shortage with the rise of TB meningitis in children. The shortage in global BCG supply has also led to a rethinking of the established treatment guidelines in rationing of the administration of BCG for bladder cancer ^20^.

Results from the Log_10_ CFU/lungs data demonstrates that the lowest dose at 30 CFU failed to protect both strains of adult mice when infected via aerosol route with either H37Rv or HN878. However, in the study conducted by Kiros *et al.*, 2010 ^17^, BALB/c mice vaccinated with BCG Danish at 30 CFU protected mice from H37Rv infection via intravenous route of challenge and the study used infant mice. For BCG doses at 3×10^5^ to 300 CFU in both strains of mice, protection against TB was influenced by the type of infecting *Mtb* strain. H37Rv infected BALB/c and F1 mice displayed significant protection in the lungs when vaccinated with BCG doses at 3×10^5^ to 300 CFU. However, for HN878 infected mice, the lowest protective BCG dose for BALB/c mice was BCG at 3000 CFU and for F1 mice it was 300 CFU. Furthermore, as per the increasing doses of BCG, a decremental trend in the TB burden (Log10 CFU/lungs) was observed in the lungs of BALB/c and F1 mice when infected with HN878 whereas, this trend was not seen for H37Rv infected BALB/c and F1 mice.

CD4 T cells play a central role for protective immunity against TB, as described by several human studies and animal models of TB ^21, 22, 23^. The important findings from Sallusto and her colleagues ^24, 25, 26^ indicated that there were two important subsets of memory T cells - TEM and central memory T cells (TCM). Differentiation of these memory T cells are based on the clear phenotypic characteristics - TEM are CD44^hi^ CD62L^lo^ CDR7^lo^ while, TCM are CD44^hi^ CD62L^hi^ CCR7^hi^. However, recent evidence depicts a complex interplay and diverse functionality under the broad subsets of memory T cells ^27^. Kaveh *et al,* 2011 ^28^ reported TEM cells (CD4+ CD44^hi^ O)62L CD27-expressing IFNγ/IL2/TNFα) persisted for 18 months following BCG vaccination in BALB/c mice which were strongly associated with the protection against *M. bovis* ^29^. When we examined TEM CD4^+^CD44^hi^CD62L^lo^ expressing IFNγ/IL2/TNFα in the spleens of BALB/c and F1 mice, the TEM cells were absent in the BCG 300 and 30 CFU groups for BALB/c mice while, in F1 mice, TEM cells were absent for BCG 30 CFU group only. The correlation of TEM cells expressing IFNγ/IL2/TNFα for BCG dose groups and the correspondent protection data (Log_10_ CFU/lungs) in the lungs of BALB/c and F1 mice, showed significant correlation for both strains of mice when infected with HN878 but, no correlation when infected with H37Rv. Interestingly, the protection data (Log_10_ CFU/lungs) in the lungs of BALB/c mice for BCG vaccinated groups significantly correlated with TEM cells expressing IFNγ/TNFα. Although, for the BCG 300 CFU group in BALB/c mice, TEM cells were not detected and yet, when infected with H37Rv, the protection was equivalent to BCG 3 x10^4^ – 3000 CFU groups whereas, BCG at 300 CFU group could not protect against HN878 infection. This is an interesting observation, if the protection in H37Rv infected BALB/c mice was not due to double or triple cytokine expressing CD4 TEM, this may suggest that either the protection was from the TEM independent mechanism or due to strain heterogeneity of *Mtb* or the presence of virtually undetected TEM. BCG at 300 CFU group in BALB/c mice is the lowest protective dose which is influenced by the type of *Mtb* infecting strain and would be an interesting dose of choice for research to understand BCG efficacy against different *Mtb* strains.

Examination of the cytokine responses in the M7 stimulated cells isolated from the lung and spleen revealed a distinct pattern of cytokine/chemokine between BALB/c and F1 mice. IFNγ is critical for the TB protection and BCG is the only vaccine whose protective immunity is believed to be depended upon the induction of CD4^+^ T cells that produce IFNγ, which in turn activates macrophages to kill *Mtb* ^30, 31^. As we expected, IFNγ responses in the supernatants of M7 stimulated splenocytes and lung cells of BCG groups showed correlation with the *ex-vivo* IFNγ responses for both strains of mice (Supplementary Fig. 1). In addition, IL2 responses albeit, low in concentration compared to the IFNγ, were detected in BCG 3 x 10^5^ −300 CFU groups. The IL2 responses from the supernatants of stimulated splenocytes were significantly inversely correlated with the *Mtb* burden in the lungs of F1 mice infected with either H37Rv or HN878. This suggests that the low concentration of detected IL2 in the spleen would be an indication for the exacerbation of *Mtb* in the lungs of F1 infected mice and can be used as an immunological marker for F1 mice in preclinical TB vaccine testing. However, in BALB/c mice, IL2 responses either from the supernatants of stimulated splenocytes or lung cells were not correlated with the *Mtb* burden in the lungs of infected mice. Interestingly, there were unexpected and differential responses of the cytokine/chemokine concentrations in the lungs of F1 mice and spleen of BALB/c mice. In the lungs of F1 and the spleen of BALB/c mice, the concentrations of IL6, TNFα, CCL4, CCL22, CCL5, CCL17 and CCL2, CXCL1, CXCL2, respectively, were higher in the control group compared to the BCG vaccinated groups. Surprisingly, the concentrations of cytokine/chemokine in the lowest dose group, 30 CFU which failed to protect *Mtb* infected BALB/c and F1 mice followed the cytokine/chemokine responses almost like BCG 300 CFU and not the control group. High dose of BCG has been implicated to skew immune responses in the mixture of Th1/Th2, but we could not detect any responses for IL4 in BALB/c or F1 mice.

This study has provided some interesting suppositions and questions: 1. Though there were no protection in the lungs of BALB/c or F1 mice after BCG vaccination with 30 CFU, we were surprised to find that the cytokine/chemokine responses were not like the control group. In fact, they were more like those in the BCG 300 CFU group, which indeed protected F1 and BALB/c mice infected with H37Rv. 2. Many animal studies commonly use BCG dose at 3 x 10^5^ CFU and have shown to persist in mice for 18 months ^28^, the protection against *Mtb* in these mice is suggested to be from constant priming of TEM by the presence of live BCG ^29^. Does this depicts that the loss of protection in the lowest dose at 30 CFU is due to the waning of BCG which subsequently ceases constant priming of TEM cells or is it possible that BCG dose at 30 CFU is so low that the frequency of TEM cells failed to establish at an optimum level to fight against *Mtb* infection? C. Depending on the research question, 300 CFU would be an interesting BCG dose to study in BALB/c and F1 mice, for example, BCG dose at 3 x 10^5^ CFU persists in organs for many months in mice and use of BCG dose at 300 CFU may shorten the experimental time and effectively provide a realistic time scale to speed up a waning experiment.

Finally, we conclude that BCG dose at 3000 CFU per mouse, a human equivalent dose, protects BALB/c and F1 mice when infected with H37Rv or HN878 which also is a realistic window to test differences in the protection of the new TB vaccine against the benchmark BCG vaccine. The combination of *Mtb* and mouse strain is an important parameter for evaluating new TB vaccine against different clinical isolates and must be considered. The spleen IL2 alongside lungs IFNγ from the supernatants of stimulated splenocytes or lung cells may be used as an important immunological marker for a preclinical TB vaccine testing in F1 mice.

## Methods

### Ethics and animals

All animal procedures were performed in accordance with the UK Home Office (Scientific Procedures) Act 1986; under appropriate licences, the study protocol was approved by the NIBSC Animal Welfare and Ethics Review Body.

BALB/c and F1 mice, female, 8 weeks old and with approximately 15 −20gm body weight were obtained from the SPF facilities at the Charles River UK Ltd. All animals were housed in an appropriate BSL2 (BCG vaccinated) and BSL3 (TB infected) containment facilities at NIBSC. Mice were checked daily throughout the experiment by trained animal technicians and this frequency could increase if any adverse reactions were observed. All mice were weighed before TB infection and subsequently weighed weekly post infection until scheduled experiment termination. No adverse effect or severe weight loss (more than 20% body weight) were observed in any mice for the entire duration of the study.

### Study groups

The study was divided into two parts: 1, H37Rv or HN878 challenge; BALB/c and F1 mice, 5 mice per group were vaccinated via intradermal route with various doses of BCG Danish 1331 and sterile saline was administered to the control group. Four weeks post vaccination, all groups (5 mice/group) were infected with *Mtb* H37Rv or HN878 via aerosol route. BALB/c and F1 mice infected with H37Rv had BCG at 3×10^5^ – 30 CFU whereas, in HN878 infected BALB/c and F1 mice, all BCG vaccinated groups except 3×10^4^ CFU were selected. 2, Immunogenicity evaluation; BALB/c and F1 mice were receiving BCG vaccination via the intradermal route with various doses except 3×10^4^ CFU. At 6 weeks post vaccination, all mice were terminated, the spleen and lungs were harvested for preparation of single cell suspension for subsequent immunological analyses.

### BCG vaccination and Mtb Challenge

BCG vaccination for BALB/c and F1 mice for H37Rv challenge experiment was performed using an expired commercial preparation of Staten Serum Institute BCG vaccine stored at −20°C. However, for HN878 challenge experiment and immunogenicity evaluation, the WHO Reference Reagent for BCG vaccine of Danish 1331 substrain preparation (NIBSC code 07/270 ^14^; https://www.nibsc.org/products/brm_product_catalogue/detail_page.aspx?catid=07/270) was used. Lyophilised BCG preparations were reconstituted and diluted in sterile saline to obtain approximately 6 x 10^6^ CFU/ml. Subsequent dilutions (Supplementary Table 1) were prepared in sterile saline to obtain various doses of 3 x 10^5^ to 30 CFU/mouse, administered in 2 × 25 μL injections by bilateral intradermal route whereas, sterile saline was administered in the control group. All BCG doses (3 x 10^5^ to 30 CFU) were checked by plating in duplicate onto 7H11 agar plates (DIFCO) supplemented with Oleic Acid Dextrose Catalase (OADC, BD) and CFU/ml were estimated at 4 weeks post incubation at 37^0^C.

For low-dose aerosol infections, H37RV or HN878 frozen stocks (in 7H9 broth containing 0.05% Tween 80, OADC and 1% glycerol) were diluted in sterile distilled water to approximately 5-7 × 10^6^ CFU/ml and placed in a glass nebulizer attached to a Nebuliser system (Walkers, UK)as described previously ^32, 33^. The mice were exposed to an aerosol infection in which live bacilli were deposited in the lungs of each mouse (expected ~100 CFU/ lungs). Confirmation of low dose infection for H37Rv and HN878 challenge experiment was performed by necropsied 4 or 5 BALB/c mice (Supplementary Table 2) on the same day of challenge. Lung lobes were removed, homogenised and serially diluted samples were plated onto 7H11 agar (DIFCO) supplemented with OADC (BD). Bacterial colonies were enumerated 3-4 weeks post incubation at 37^0^C. Upon termination of the experiment at 4 weeks post challenge, mice were necropsied, the spleen and lungs were removed, homogenized and serially diluted samples were plated in duplicate onto OADC-supplemented 7H11 agar plates. The plates were incubated at 37 °C and bacterial colonies were enumerated after 3-4 weeks.

### Splenocytes and lung cells preparations

For the immunogenicity evaluation part; the spleen and lungs were harvested for isolating single cell suspension. For splenocytes isolation, the spleen was mashed through a 40□μm cell strainer, washed at 300 g for 5 min and resuspended at 1□×□10^7^ cells/ml in the supplemented Dulbecco’s Modified Eagle’s Medium (DMEM) (Sigma) containing 10% heat inactivated foetal calf serum (Biosera, UK) and 2% penicillin/streptomycin (Gibco) for use in subsequent assays.

Harvested lung lobes were minced and agitated for 1 □hr at 37 °C in supplemented DMEM medium containing 50 U/ml collagenase I (Gibco) and 10 U/ml DNase II (Sigma), then followed by passing through a 40□μm cell strainer. Isolated lung cells were washed and resuspended in supplemented DMEM at 1□×□10^7^ cells/ml for use in subsequent assays.

### Mycobacterial antigens (M7 protein cocktail)

The M7 protein cocktail, which is made of a pool of 7 immunogenic recombinant mycobacterial proteins - Rv1886c (Ag85B), Rv0251c (hsp20), Rv0287 (TB9.8), Rv0288 (TB10.4), Rv3019c (TB10.3), Rv3763, Rv3804c (Ag85A) (Lionex, GmbH, Germany) and stored at −80^0^C, was used in assays where cells required *in vitro* stimulation. Each protein concentration in the cocktail is 50 μg/ ml and for stimulating cells, M7 cocktail was diluted to achieve a final concentration of 2 μg/ ml for each protein. M7 protein cocktail can be purchased from the NIBSC website.

### ELISPOT

Approximately 2×10^5^ splenocytes or lung cells were incubated in duplicate in 96-well filter plates (MSIPS4510 Millipore, Ireland) with or without M7 protein cocktail (final conc. 2 μg/ ml), or concanavalin A (10 μg/ml, Sigma) as a positive control for approximately 16 h at 37°C. The frequency of IFNγ secretors was detected by the AID ISPOT (ELR08IFL) reader (AID Autoimmun Diagnostika, Germany), as per manufacturer’s instructions.

### Flow Cytometry for surface markers and intracellular cytokines

Splenocytes were cultured with 2 □ μg/ml of M7 protein cocktail, 1 μg/ml of anti-CD28 (BD Biosciences) for 2 hrs at 37□°C under 5% CO_2_, after which 10 μg/ml Brefeldin A (Sigma) was added for a further incubation of 14 hrs. Subsequently, cells were washed at 300□g for 5 min and surface stained with a combination of pre-titrated monoclonal antibody conjugates: CD90.2 - eFluor 450; CD27 - PerCP-eFluor710 (both Life Technologies, UK); CD62L - FITC; CD8 - Alexa Fluor 700; CD44 - Brilliant Violet 785; Zombie Aqua™ Fixable Viability Dye (all BioLegend) and CD4 - APC-H7 (BD Biosciences). Cells were then washed, fixed, permeabilised and stained intracellularly using BD Cytofix and BD Cytoperm (BD Bioscience), as per manufacturer’s instructions, with a combination of: IL17a – PE; IFNγ - PE-Cy7; IL2 - APC; and TNFα-BV605 (all BioLegend). Cells were analysed immediately after final staining. Data were acquired using a SORP LSR Fortessa (BD Bioscience), utilizing a 532 nm laser for PE and PE-Cy7, and analysed on Flowjo v.10.1 software (BD Biosciences). All analyses were gated on a minimum of 100,000 live lymphocytes. Compensation was performed using UltraComp eBeads (Life Technologies) according to the manufacturer’s instructions. Fluorescence minus one control were used to set gates for cytokine analyses.

### Multiplex cytokine/chemokine bead assays

Splenocytes and lung cells at ~ 2 x 10^6^ cells/ ml were stimulated with or without M7 cocktail (final conc. 2 μg/ ml) and incubated at 37^0^C under 5% CO_2_ for 3 days. Supernatants were harvested and cytokine/ chemokine beads assays were performed using BioRad’s Bio-Plex pro mouse cytokine 34 or 31 - plex assay. BioRad’s 34-plex was used for experiments using BALB/c mice and, due to the availability of the reagent kit, 31-Plex was used for experiments using F1 mice (Table 2). All multiplex magnetic bead assays were performed according to the manufacturer’s protocol. Concentrations of cytokine/chemokine were extrapolated using manufacturer’s provided standard cocktail in the kits. Reported concentrations of cytokine/chemokine (Supplementary Table 3 and Fig. 4A&B) were obtained from the M7 stimulated cells subtracted by the unstimulated cells.

### Statistical analyses

All data were analysed with the GraphPad Prism 8 software (Graph Pad, USA) using one-or two-way ANOVA with Dunnett’s or Tukey’s post-test (3-treatment groups), respectively. Mycobacterial CFUs were Log10 transferred before comparison. Correlation coefficients were assessed using Pearson’s two-tailed correlation test with 95% confidence interval. Differences with a ρ value <0.05 were considered and denoted with, *p<0.05, **p<0.005, ***p<0.0005, ****p<0.0001.

## Acknowledgements

This work is partly funded by the European Commission under grant agreement No. 730964 TRANSVAC2 project. We extend our thank you to all staff at Biological Services Unit for their continuous effort in supporting our animal research with the highest standards of animal care and husbandry. We thank TB CL3 manager Sara Goulding and Vicky Rannow for smooth operation of the facility and to uphold safe working practices in a TB high containment facility.

## Author Contributions

B.K. contributed to the experiments, study design, interpretation, analyses and manuscript preparation. J.K. and B.D. contributed to the experiments and study design. D.A.K contributed in providing M7 protein cocktail, performing flow cytometry analyses and interpretation. P.J.H contributed to the study design. M.M.H contributed to the study design and extensive manuscript revision. All authors approved the final version of the manuscript and have agreed to be personally accountable for their respective contributions and ensure that questions related to the accuracy or integrity of any part of the work, even those in which the author was not personally involved, are appropriately investigated, resolved, and the resolution documented in the literature.

## Competing interests

The authors declare no competing interests.

## Additional information

Appended supplementary information as a separate word document - supplementary information.docx

## Notes

### Competing Interest Statement

The authors have declared no competing interest.

