## Supplementary information for "Efficacy and immunogenicity of different BCG doses in BALB/c and CB6F1 mice when challenged with H37Rv or Beijing HN878"

#### Supplementary Table 1: Enumeration of BCG CFU/ml

| Expected BCG CFU/ml | SSI BCG Danish 1331 strain CFU/ml for H37Rv challenge experiment | | WHO RR BCG Danish 1331 strain (NIBSC code 07/270) CFU/ml for HN878 challenge experiment | |
|---|---|---|---|---|
| | **Average CFU/ml enumeration using duplicate 7H11 plates** | **Actual dose of BCG CFU per mouse (50µl given by ID route)** | **Average CFU/ml enumeration using duplicate 7H11 plates** | **Actual dose of BCG CFU per mouse (50µl given by ID route)** |
| **$6x10^6$ CFU/ml** | $4.3 \times 10^6$ CFU/ml | $2.15 \times 10^5$ CFU | $2.4 \times 10^6$ CFU/ml | $1.2 \times 10^5$ CFU |
| **$6x10^5$ CFU/ml** | $3.75 \times 10^5$ CFU/ml | $1.88 \times 10^4$ CFU | Excluded from the exp. | Excluded from the exp. |
| **$6x10^4$ CFU/ml** | $3.85 \times 10^4$ CFU/ml | 1930 CFU | $2 \times 10^4$ CFU/ml | 1000 CFU |
| **6000 CFU/ml** | 2200 CFU/ml | 110 CFU | 3000 CFU/ml | 150 CFU |
| **60 CFU/ml** | 24 CFU/ml | 12 CFU | 34 CFU/ml | 17 CFU |

#### Supplementary Table 2: Enumeration of CFU/ml for Infection inoculum and the lungs of mice on the same day of aerosol infection with H37Rv and HN878.

| | **H37Rv challenge** | **HN878 challenge** |
|---|---|---|
| **Expected challenge inoculum** | $5.45 \times 10^6$ CFU/ml | $7 \times 10^6$ CFU/ml |
| **Average CFU/ml enumeration using duplicate 7H11 plate** | $3.6\text{-}5 \times 10^6$ CFU/ml | $4\text{-}7 \times 10^6$ CFU/ml |
| **Average CFU in the lungs of five BALB/c mice (Av. CFU ± SD)** | 71 ± 64 CFU | 144 ± 63 CFU |

### Supplementary Table 3: Cytokine/chemokine values in pg/ml using Bio-plex Manager 6.1 software.

| Cytokine/Chemokine | BALB/c - Spleen | | | | | F1 - Spleen | | | | | BALB/c - Lungs | | | | | F1 - Lungs | | | | |
|---|---|---|---|---|---|---|---|---|---|---|---|---|---|---|---|---|---|---|---|---|
| | Control | 3.E+05 | 3000 | 300 | 30 | Control | 3.E+05 | 3000 | 300 | 30 | Control | 3.E+05 | 3000 | 300 | 30 | Control | 3.E+05 | 3000 | 300 | 30 |
| CXCL13 | 168.87 | 170.69 | 150.70 | 24.17 | 105.32 | 69.53 | 41.98 | 35.11 | 53.40 | 84.86 | 121.87 | 114.40 | 132.95 | 48.17 | 151.29 | 25.81 | -1.81 | 47.28 | 33.47 | 8.96 |
| CCL27 | 153.97 | 182.53 | 115.66 | 23.64 | -13.98 | 119.39 | 134.62 | 57.62 | 193.54 | 160.54 | 72.36 | 153.76 | 144.39 | -6.37 | 113.67 | 331.29 | 243.39 | 261.43 | 428.30 | 69.39 |
| CXCL5 | 113.85 | 119.82 | 100.28 | 3.52 | 28.12 | 38.53 | 39.27 | 28.04 | 57.05 | 60.51 | 207.05 | 937.72 | 194.78 | 121.61 | 70.65 | 579.98 | 463.82 | 439.14 | 456.82 | 303.55 |
| CCL11 | 2.83 | 2.63 | 1.93 | 0.00 | 0.00 | 1.27 | 1.05 | 0.43 | 1.33 | 1.34 | 2.66 | 4.41 | 2.97 | 0.67 | 1.51 | 2.28 | 2.63 | 1.89 | 2.52 | 1.21 |
| CCL24 | 0.00 | 0.00 | 0.00 | 0.00 | 0.00 | 0.00 | 0.00 | 0.00 | 0.00 | 0.00 | -267.46 | -248.19 | -432.30 | -157.59 | 8.63 | -741.23 | -449.21 | -370.47 | -313.01 | -344.96 |
| CX3CL1 | 5.05 | 5.05 | 2.90 | -24.27 | -41.30 | 7.72 | 18.22 | -0.53 | 8.76 | 13.38 | 4.07 | 13.02 | 9.78 | -3.65 | 0.00 | 6.93 | 6.26 | 7.98 | 8.26 | 1.49 |
| GM-CSF | 8.26 | 8.27 | 5.70 | -26.10 | -44.58 | 7.02 | 14.44 | 12.19 | 13.57 | 8.14 | 18.54 | 47.75 | 15.48 | 8.33 | 7.89 | 60.10 | 50.55 | 43.36 | 37.39 | 29.06 |
| CCL1 | 0.95 | 27.32 | 13.36 | -26.82 | -46.05 | 1.00 | 9.91 | 5.23 | 11.82 | 2.30 | 1.41 | 129.23 | 21.17 | 0.10 | 0.42 | 2.15 | 32.21 | 14.79 | 30.26 | 1.40 |
| IFNγ | 6.58 | 728.58 | 689.73 | 5.89 | 6.32 | 99.98 | 590.08 | 362.33 | 557.85 | 121.91 | 2.75 | 85.89 | 9.99 | -0.30 | 2.40 | 2.59 | 15.92 | 6.68 | 8.94 | 1.27 |
| IL1β | 2.83 | 5.75 | 5.75 | -28.63 | -19.34 | 11.02 | 24.72 | 13.81 | 13.81 | 9.00 | 11.56 | 30.39 | 12.83 | 0.00 | 5.13 | 6.87 | 12.87 | 8.91 | 18.43 | 10.73 |
| IL2 | 1.62 | 43.10 | 25.44 | -26.46 | -45.74 | 0.56 | 16.99 | 10.81 | 8.80 | 1.01 | 0.92 | 34.06 | 6.22 | 0.43 | 0.23 | 0.94 | 13.28 | 3.76 | 3.68 | 0.29 |
| IL4 | 0.31 | 0.94 | 0.84 | -27.09 | -45.89 | 3.50 | 0.08 | -3.90 | 2.61 | 3.04 | 0.86 | 1.38 | 0.93 | 0.33 | 0.69 | 1.91 | 1.24 | 1.90 | 4.44 | -0.99 |
| IL6 | 387.64 | 919.72 | 596.51 | 84.30 | 169.00 | 1029.57 | 1631.93 | 1410.87 | 1815.13 | 1466.01 | 1140.35 | 8510.27 | 1400.92 | 492.99 | 466.87 | 1783.44 | 1392.55 | 918.52 | 1180.71 | 601.25 |
| IL10 | 59.19 | 92.71 | 78.71 | -19.40 | -6.10 | 150.47 | 187.87 | 138.24 | 196.91 | 202.62 | 39.98 | 72.35 | 50.69 | 1.79 | 28.33 | 0.00 | 0.00 | 0.00 | 0.00 | 0.00 |
| IL16 | -197.70 | -177.85 | -102.25 | -66.36 | 209.92 | 37.42 | 24.81 | -105.50 | -99.39 | -48.73 | -31.22 | 50.44 | -32.26 | -37.25 | -4.06 | 56.42 | 72.80 | 107.21 | 97.58 | 100.09 |
| IP10 | 39.13 | 54.81 | 56.82 | -9.62 | 28.46 | 274.26 | 470.84 | 301.07 | 694.21 | 396.45 | 37.11 | 174.33 | 29.61 | -6.00 | 61.84 | 235.90 | 321.09 | 222.40 | 265.10 | 146.99 |
| CXCL11 | 31.17 | 28.13 | 22.89 | -20.59 | -24.73 | 84.58 | 112.96 | 80.61 | 139.18 | 130.62 | 29.37 | 349.88 | 44.17 | 5.09 | 26.47 | 256.82 | 196.65 | 207.51 | 273.50 | 72.12 |
| CXCL1 | 1436.42 | 1098.90 | 705.91 | 202.45 | 313.63 | 788.70 | 790.92 | 609.16 | 812.23 | 994.41 | 5523.74 | 6022.01 | 7135.67 | 2271.40 | 3078.96 | 13393.78 | 7275.09 | 7233.32 | 9781.89 | 4663.81 |
| CCL2 | 200.71 | 87.40 | 66.79 | -21.94 | -34.80 | 40.57 | 66.87 | 56.49 | 63.71 | 64.64 | 973.77 | 9086.06 | 980.94 | 642.08 | 359.00 | 1935.26 | 1739.76 | 1620.55 | 1584.96 | 931.70 |
| CCL8 | 0.00 | 0.00 | 0.00 | 0.00 | 0.00 | not included in the multiplex | | | | | 0.00 | 0.00 | 0.00 | 0.00 | 0.00 | not included in the multiplex | | | | |
| CCL7 | 45.90 | 14.05 | 12.03 | -25.06 | -41.80 | 3.88 | 5.09 | 4.62 | 6.44 | 7.31 | 110.95 | 1190.52 | 244.05 | 34.89 | 65.62 | 184.52 | 141.83 | 127.11 | 197.90 | 71.81 |
| CCL12 | 0.00 | 0.00 | 0.00 | 0.00 | 0.00 | 0.00 | 0.00 | 0.00 | 0.00 | 0.00 | 0.00 | 0.00 | 0.00 | 0.00 | 0.00 | 0.00 | 0.00 | 0.00 | 0.00 | 0.00 |
| CCL22 | 537.69 | 611.09 | 469.72 | 110.87 | 277.64 | 135.22 | 235.51 | 177.95 | 345.22 | 321.69 | 588.50 | 5473.11 | 732.04 | 263.57 | 279.16 | 392.31 | 206.03 | 153.68 | 196.50 | 137.65 |
| CCL3 | 1837.50 | 1977.79 | 1098.59 | 372.64 | 575.69 | 2264.52 | 1692.59 | 1747.28 | 2735.61 | 4499.99 | 7015.16 | 7784.34 | 7322.46 | 2586.61 | 5050.60 | 1334.41 | 1719.81 | 1869.16 | 4748.09 | 1195.80 |
| CCL4 | 893.56 | 977.03 | 638.52 | 112.99 | 260.74 | 628.36 | 808.47 | 696.04 | 877.43 | 799.87 | 1144.69 | 2511.22 | 1738.81 | 547.90 | 632.20 | 1739.16 | 1161.62 | 784.57 | 933.29 | 546.65 |
| CXCL2 | 4265.26 | 3790.41 | 2301.22 | 722.75 | 1154.43 | not included in the multiplex | | | | | 56136.94 | 293922.42 | 627063.28 | 8264.20 | 17789.77 | not included in the multiplex | | | | |
| CCL20 | 0.00 | 0.00 | 0.00 | 0.00 | 0.00 | 0.00 | 0.00 | 0.00 | 0.00 | 0.00 | -2.73 | 0.75 | -0.02 | -1.84 | 1.22 | 0.00 | 0.00 | 0.00 | 0.00 | 0.00 |
| CCL19 | 8.75 | 13.93 | 13.11 | -23.60 | -38.04 | 17.10 | 26.06 | 15.23 | 28.89 | 36.90 | 6.48 | 11.26 | 2.92 | -0.66 | 10.33 | 27.32 | 22.77 | -3.26 | 35.77 | 14.80 |
| CCL5 | 146.31 | 420.80 | 283.45 | 41.33 | 97.91 | 495.87 | 653.70 | 514.05 | 630.39 | 531.09 | 262.78 | 1237.95 | 347.81 | 157.65 | 202.02 | 801.52 | 654.37 | 515.44 | 643.18 | 315.44 |
| CXCL16 | 6.17 | 6.53 | 5.30 | 0.00 | OOR< | 6.65 | 6.08 | 4.00 | 5.02 | 7.22 | 24.39 | 161.01 | 40.44 | 5.53 | 11.90 | 48.66 | 32.77 | 30.15 | 30.49 | 14.51 |
| CXCL12 | 23.71 | 27.87 | 26.51 | -20.35 | -20.92 | 87.47 | 86.05 | 82.06 | 99.65 | 118.72 | 33.88 | 49.16 | 25.11 | 10.30 | 29.75 | 76.90 | 81.14 | 61.46 | 77.82 | 53.91 |
| CCL17 | 8.97 | 9.22 | 7.24 | -25.09 | -42.90 | 13.49 | 12.64 | 8.51 | 16.35 | 15.39 | 17.89 | 73.16 | 12.23 | 1.17 | 8.03 | 54.45 | 31.00 | 28.31 | 40.99 | 21.76 |
| TNFα | 32.93 | 23.15 | 9.93 | -25.60 | -42.25 | 11.59 | 16.48 | 7.60 | 12.01 | 10.51 | 94.35 | 510.31 | 139.07 | 32.61 | 23.54 | 26.56 | 19.87 | 13.48 | 12.91 | 7.90 |
| CCL25 | 0.00 | 0.00 | 0.00 | 0.00 | 0.00 | not included in the multiplex | | | | | 0.00 | 0.00 | 0.00 | 0.00 | 0.00 | not included in the multiplex | | | | |

Note: All concentrations in pg/ml were obtained from the linear part of the 5P standard curve using Bio-Plex Manager 6.1 software. The concentrations (pg/ml) in the table represent the result after subtracting concentrations of cytokines/chemokines in the supernatant of M7 stimulated cells with those from unstimulated cells. Blue highlighted cells in the table shows significance (p≤0.05) when compared to the control group.

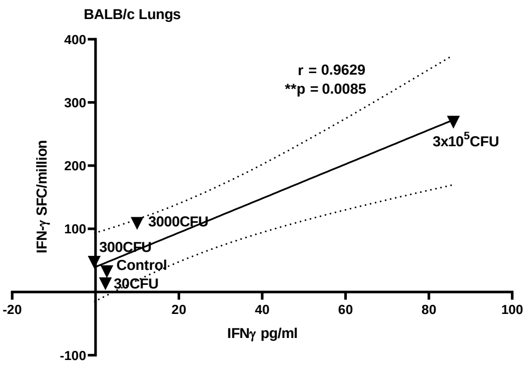


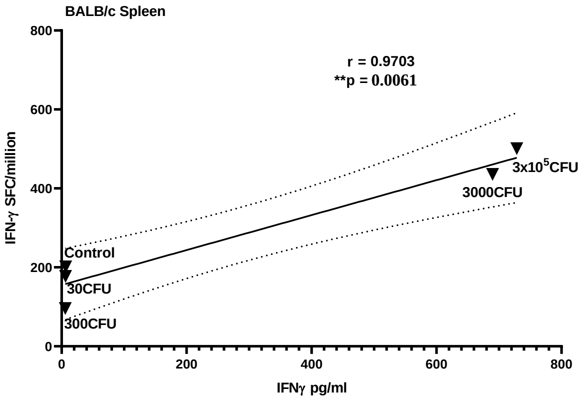


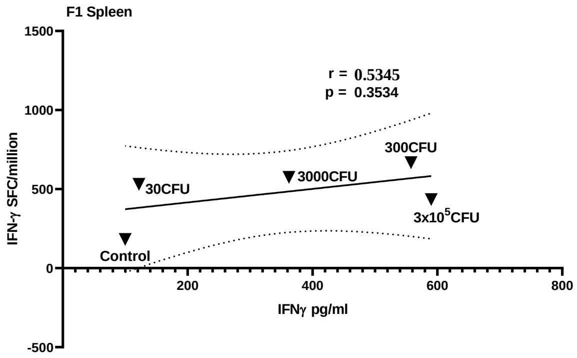


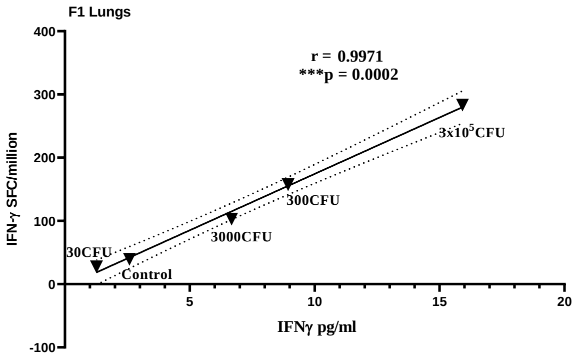

**Supplementary Figure 1: Correlation of *ex-vivo* IFNγ release from cells with the observed concentration of IFNγ in the supernatants of stimulated cells.** Representative graphs for correlation analysis between the mean of IFNγ cytokine/chemokine from supernatants of stimulated splenocytes/lung cells and ex-vivo IFNγ (SFC/million) in the lungs of BCG vaccinated F1 and BALB/c mice. Correlation coefficients were assessed using Pearson's two-tailed correlation test with 95% confidence interval shown as dotted lines in A and B. The control and BCG doses in vaccinated groups have been annotated on the graphs to highlight the correspondent data.

**Supplementary Table 4: Other cytokines/ chemokines showing statistically significant correlations with IFNγ in the supernatants of M7 stimulated splenocytes and lung cells from mice vaccinated with BCG vaccine**.

| | BALB/c Splenocytes IFNγ correlation with other cytokines | | F1 splenocytes - IFNγ correlation with other cytokines | | | BALB/c lung cells IFNγ correlation with other cytokines | F1 lung cells IFNγ correlation with other cytokines |
|---|---|---|---|---|---|---|---|
| | IL6 | CCL5 | IL2 | CCL1 | CCL5 | IL2 | IL2 |
| **Pearson r** | 0.7998 | 0.9280 | 0.8114 | 0.9701 | 0.8883 | 0.9970 | 0.9258 |
| **P value (2-tailed)** | 0.0406 | 0.0229 | 0.0370 | 0.0062 | 0.0441 | 0.0002 | 0.0088 |
| **Significance** | * | * | * | ** | * | *** | ** |
| **95% confidence intervals (Slope)** | 0.062 to 1.50 | 0.097 to 0.642 | 0.003 to 0.050 | 0.010 to 0.028 | 0.013 to 0.536 | 0.336 to 0.449 | 0.413 to 1.309 |
